# The Splicing Factor PTBP1 interacts with RUNX1 and is Required for Leukemia Cell Survival

**DOI:** 10.1101/2025.05.16.654547

**Authors:** Arjun Dhir, Alexander Ethell, Riley Watkins, Calvin Lam, Kevin Tur-Rodriguez, Jimmie Persinger, Kasidy K. Dobish, Sipra Panda, Shannon M. Buckley, Samantha A Swenson, Sandipan Brahma, M. Jordan Rowley, R. Katherine Hyde

## Abstract

Runt-related Transcription Factor 1 (RUNX1) is essential for definitive hematopoiesis and is among the most frequently mutated genes in leukemia. Previous work from our lab demonstrated that Histone Deacetylase 1 (HDAC1), a known RUNX1 partner, is unexpectedly required for active transcription suggesting a non-histone role for HDAC1 regulating components of the RUNX1 complex. Here, we use proteomics, genomics, and long-read transcriptomics to identify novel RUNX1 interacting partners and decipher their role in gene regulation and RNA splicing in leukemia cells. We demonstrate that Polypyrimidine Tract Binding Protein 1 (PTBP1) interacts with RUNX1 in an HDAC1 dependent manner. Chromatin profiling revealed extensive genome-wide overlap in sites occupied by RUNX1 and PTBP1, with significant enrichment at promoters of actively transcribed genes. Loss of PTBP1 in AML cells led to widespread alterations in RNA splicing and decreased expression of genes whose promoters are bound by both factors, including metabolic genes. In agreement with these findings, we found that loss of PTBP1 reduced glycolysis and glucose uptake and ultimately caused cell death. Based on our data, we propose that the interaction between RUNX1 and PTBP1 facilitates expression of metabolic proteins essential for leukemia cell growth and survival.

**Graphical Abstract:** 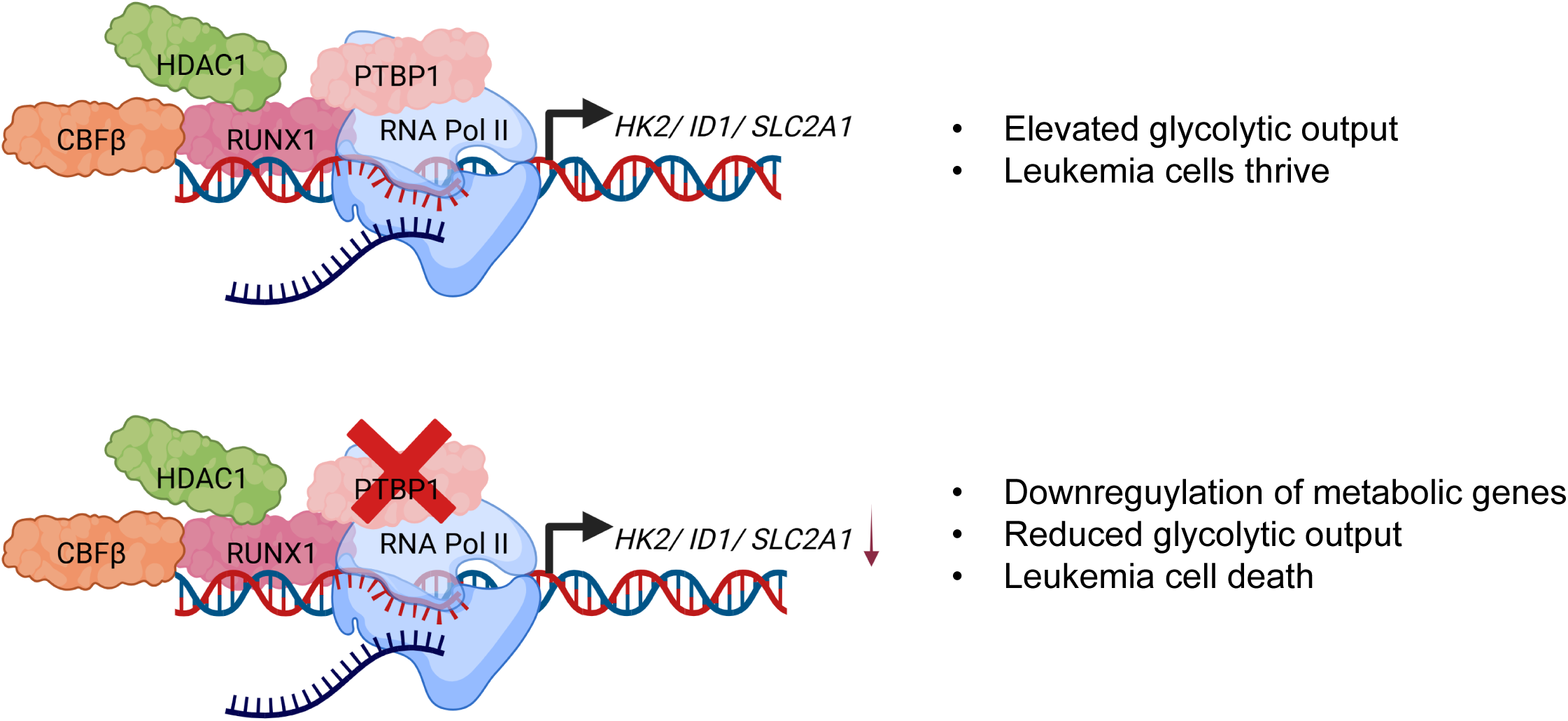

**KEY POINTS:**

- PTBP1 binds RUNX1 in a HDAC1-dependent manner and co-localizes to the promoters of target genes in leukemia cells.
- Loss of PTBP1 decreases expression of key metabolic genes, resulting in decreased cell growth and glycolysis, increased sensitivity to chemotherapy, and cell death.

## INTRODUCTION

Leukemia is a cancer affecting blood and bone marrow cells, often caused by mutations involving key hematopoietic transcription factors, such as RUNX1. Germline mutations in RUNX1 are associated with the leukemia predisposition syndrome Familial Platelet Disorder with Associated Myeloid Malignancies (1). Somatic deletions, point mutations, and oncogenic fusion proteins involving RUNX1 (RUNX1::RUNX1T1, ETV6::RUNX1) and its obligate binding partner CBFβ (CBFβ::SMMHC) are common in Acute Myeloid Leukemia (AML) and Acute Lymphoblastic Leukemia (ALL), making RUNX1 one of the most frequently mutated genes in leukemia (2, 3).

RUNX1 (Runt-related transcription factor 1, AML1) is a member of the core binding factor (CBF) family of proteins, which are known to have roles in embryonic development, proliferation, differentiation, and cancer (4–6). RUNX1 is known to interact with several other transcription factors and epigenetic modifiers, including Histone Deacetylase 1 (HDAC1) (7). HDAC1 is best known for its role in transcriptional repression but can also mediate protein interactions and promote active transcription (8). Our lab previously showed that HDAC1 is required for active transcription of RUNX1 target genes in a mouse model of CBFβ::SMMHC (CM) driven AML, making leukemia cells particularly sensitive to the HDAC1 inhibitor, entinostat (9). This finding raised the possibility that HDAC1 activity may be required to mediate interactions between RUNX1 and other co-factors to promote target gene expression.

In this study, we identified a new RUNX1 binding partner, the splicing regulator, Polypyrimidine Tract Binding Protein 1 (PTBP1). As its name suggests, PTBP1 binds polypyrimidine-rich RNA. It is involved in pre-mRNA processing, RNA transport, alternative splicing, and is expressed nearly ubiquitously (10–13). PTBP1 has been implicated in the pathogenesis of solid tumors, but PTBP1’s role in the hematopoietic compartment has been less studied (14, 15). Knockout mouse studies indicate that PTBP1 regulates hematopoietic stem cell maintenance, erythroid differentiation, and B-cell selection (16, 17). In leukemia, PTBP1 has been linked to leukemia cell survival, metabolism, and progression (18) (19–22). However, the mechanisms behind how PTBP1 orchestrates these functions remains largely unknown. In this current study, we show that RUNX1 interacts with PTBP1 in both AML and ALL cells, and that this interaction requires HDAC1 activity. We also show that RUNX1 and PTBP1 co-localize at the promoters of actively transcribed genes including key metabolic genes, and that loss of PTBP1 decreases expression of target genes resulting in impaired glycolysis and cell death, linking RUNX1 to the control of metabolism in leukemia.

## METHODS

### Mice

All animal experiments were approved by the University of Nebraska Medical Center IACUC in accordance with NIH guidelines. *Cbfb^+/56M^*, *Mx1-Cre^+^* (*CM^+^*) or Cbfb*^+/56M^*, *Mx1-Cre^+^*, *Gt(ROSA)26Sortm4(ACTB-tdTomato, -EGFP/Luo/J)* (*Rosa^26tdT/GFP^*) (*CM^+^*, *GFP^+^*) were maintained on a mixed background (C57BL6/129S) (23, 24). To induce leukemia, mice (6-8 weeks) were treated with Polyinosine-polycytidylic acid (pI:pC) (InvivoGen, San Diego, CA), intraperitoneally (250µg/dose, three times a week, every other week for 6 weeks). Transplantations to expand leukemia cells were performed by injecting 1 x 10^5^ cells retro-orbitally into sub-lethally irradiated congenic mice, as previously described (9, 25).

### Cell Culture

*Cbfb^+/56M^, Mx1-Cre^+^* cells were cultured in StemSpan SFEM I (Stemcell Technologies, Vancouver, Canada), supplemented with 1% Lipid Mixture (Sigma-Aldrich, St. Louis, MO), 1% penicillin/streptomycin/L-glutamine (PSL), recombinant murine IL-3 (10ng/mL), IL-6 (10ng/mL) and SCF (20ng/mL) (Gibco, Grand Island, NY). Entinostat was procured from Cayman Chemical (Ann Arbor, MI). ME-1 cells (a kind gift from P. Liu, NGHRI, NIH). Kasumi-1, MV-4-11, REH, MOLT-4, and Jurkat cells were obtained from ATCC (Manassa, VA). SEM and MOLM-13 cells were obtained from DSMZ (Leibniz Institute, Germany). Mobilized CD34+ cells from healthy donors were obtained from Fred Hutchinson Cancer Center Co-operative Center for Excellence in Hematology. Patient AML cells were obtained from the Stem Cell and Xenograft Core at the University of Pennsylvania Perelman School of Medicine and Cureline Translation CRO and cultured as previously described (26).

### Colony forming assay

Colony forming assays were performed using MethoCult GF M3434 (StemCell Technologies). *CM^+^, GFP^+^* were sorted to isolate leukemic cells. 100,000 *CM^+^*, *GFP*^+^ cells were plated in 500 µl of MethoCult/well of a 24 well plate, in triplicate and counted 10 days later.

### Plasmid Constructs

The Myc tag WT PTBP1 construct was purchased from Addgene (Plasmid# 23024). The FLAG-RUNX1 WT and FLAG-RUNX1-R232, FLAG-RUNX1-Y287 and FLAG-RUNX1-G365R constructs were a kind gift from J. Yang, St. Jude’s (27). The WT-RUNX1 and RUNX1-Y407 constructs were generously shared by Paul Liu at NIH (28, 29). The RUNX1-R320 construct was a kind gift from Dong-Er Zhang at the University of California, San Diego (30).

### Site Directed Mutagenesis

Myc tag WT PTB (Addgene) was mutated to Myc-PTB-RRM1 (Y193) using the Phusion Hot Start II High-Fidelity PCR Master Mix (Thermo, Waltham, MA). Primer sequences used to generate mutant constructs are available upon request.

### Lentiviral mediated shRNA knockdown

HEK293T cells were co-transfected with second generation lentiviral plasmids, pPax2 and MD2.G (5µg each) and 10µg of the PTBP1 or Scrambled (SCR) shRNA plasmid to generate lentivirus (22). The PTBP1 knockdown and control plasmids, which express the shRNAs and the ΔLNGFR receptor under the control of a doxycycline inducible promoter were modified by GenScript (Piscataway, New Jersey), replacing puromycin resistance gene for mCherry. Virus was concentrated using Amicon® Ultra-15 Centrifugal Filter Units (MilliporeSigma, Burlington, MA). Transduction protocol for leukemia cells is available on request.

### Immunoprecipitation and Western blot

Protein overexpression studies were carried out in HEK293T cells using lipofectamine-2000 or lipofectamine-3000 (Thermo). Protein lysates were incubated with primary antibodies followed by Protein A/G Dynabeads (Thermo) or withanti-FLAG M2 magnetic beads (Sigma-Aldrich). Antibody list, applications, and dilutions used are available in Supplemental Table 1.

### Mass Spectrometry

Mouse *CM*^+^ cells incubated with entinostat or DMSO for 24 hours and nuclear lysates were prepared and immunoprecipitated with an anti-RUNX1 antibody (Abcam, Cambridge, UK) and analyzed by the UNMC Mass Spectrometry Core Facility. Quantitative data analysis was performed using progenesis QI proteomics 4.2 (Nonlinear Dynamics). Statistical analysis was performed using ANOVA with the Benjamini-Hochberg (BH) correction for multiple-testing caused false discovery rate. The adjusted p ≤ 0.05 was considered significant.

### Confocal Microscopy

Lineage depletion was performed using the EasySep™ Mouse Hematopoietic Progenitor Cell Isolation Kit (StemCell Technologies). Proximity Ligation Assay (PLA) was performed using Sigma-Aldrich’s Duolink PLA kit. Slides for both IF and PLA were imaged on the AiryScan 800 (63x magnification) at the UNMC Advanced Microscopy Facility. Quantification of PLA was carried out using the Image J plug-in, Andy’s Algorithms (31).

### Flow cytometry and cell sorting

Cells were stained with Annexin V, DAPI, CD15, CD11b, and CD271 (ΔLNGFR) (BD Biosciences, Franklin Lakes, NJ, and Biolegend, San Diego, CA) and analyzed on a BD LSRII. Cell sorting was performed on FACSAria I or FACSAria II (BDBiosciences) or BigFoot (ThermoFisher Scientific). Data was analyzed in FlowJo v.10.0.8 (FlowJo, LLC, Ashland, OR). Further details regarding antibodies can be found in Supplemental table 1.

### Cleavage Under Targets and Tagmentation (CUT&Tag)

CUT&Tag was performed as previously described (32). Primary antibodies were used at the following dilutions: PTBP1 (Thermo): 1:50; for RUNX1 (Abcam): 1:20; rabbit/mouse IgG (CST, Danvers, MA): 1:100, and H3K27me3 (CST): 1:100. For secondary antibodies (1:100), we used guinea pig anti-rabbit antibody (Antibodies Online, Pottstown, PA) and rabbit anti-mouse antibody (Abcam). Library quality and concentrations were determined using the D1000 TapeStation system (Agilent). Libraries were sequenced in 150-bp paired-end mode on the Illumina NextSeq 550 (UNMC Genomics core) or NovaSeqX 10B (Novogene) platforms, and data were analyzed as previously described (33). Adapters were clipped and paired-end reads were mapped to UCSC Hg38 using Bowtie2 (34) with parameters: --end-to-end --very-sensitive --no-mixed --no- discordant -q --phred33 -I 10 -X 700. Spike-in reads were mapped to *E. coli* K-12 genome with parameters: --end-to-end --very-sensitive --no-overlap --no-dovetail --no-unal --no-mixed --no- discordant -q --phred33 -I 10 -X 700. Continuous-valued data tracks (bedGraph and bigWig) were generated using genomecov in bedtools v2.30.0 (-bg option) and calibrated using total number of spike-in reads. Aligned reads were filtered for mapq>30, deduplicated, underwent peak identification using MACS3 (35) q-value threshold 10^-7^, and differential peaks identified using MAnorm (36).

### Long-read RNA sequencing

RNA extraction was carried out using Qiagen’s RNeasy Mini Kit (Qiagen, Germantown, Maryland) following manufacturer’s instructions. RNA was isolated from PTBP1 knockdown and control MOLM-13 cells after 9 days of doxycycline treatment. Long-read RNA sequencing was performed by CD Genomics, Shirley, NY using cDNA libraries prepared using the ONT SQK-PCS109 + SQK-PBK004 kits and sequenced on the PromethION platform. Reads were mapped to the Hg38 genome build using minimap2 (37), and filtered with a long-read mapq>50. Differentially expressed genes were identified using Stringtie (38) to generate transcript counts, followed by DESeq2 with an adj. p-value <.05. Differential isoform usage was performed by IsoformSwitchAnalyzeR (39) with a q-value >.05. Functional categorization and overrepresentation analysis was performed using EnrichR (40).

### Gene expression data

Publicly available PTBP1 gene expression data was accessed from “Normal hematopoiesis with AMLs” data set in BloodSpot 3.0, a specialized gene and protein expression database of normal and malignant hematopoietic cells (41).

### Metabolic assays

Lactate release and glucose uptake assays were performed using the Lactate Assay kit and the Glucose Uptake Assay Kit – Green (Dojindo, Rockville, Maryland).

### Statistics

Data was analyzed using either the Student’s t-test or ANOVA with Turkey post-hoc, as indicated in the figure legend, using GraphPad Prism 7. P-values ≤ 0.05 were considered significant.

## RESULTS

### PTBP1 binds RUNX1 in a HDAC1 dependent manner in leukemia

Work from our lab showed that HDAC1 binds CBFβ::SMMHC (CM) and is required for active transcription of RUNX1/CM target genes (9). Canonically, HDAC1 is associated with transcriptional repression, but can also regulate the formation of protein complexes that promote gene expression (8). We performed mass spectrometry to determine if HDAC1 activity influences the recruitment of proteins to the RUNX1 complex in leukemia cells. Five independent mouse *CM*^+^ leukemia samples were treated with 1µM entinostat, an HDAC inhibitor, or DMSO for 24 hours, nuclear lysates prepared, RUNX1 immunoprecipitated, and the precipitate analyzed by mass spectrometry (Fig 1A). After filtering for known non-specific contaminants, we identified 809 peptides, that mapped to 284 different proteins, 80 of which were identified with at least 3 unique peptides. From this list, one protein showed a statistically significant difference between control and entinostat treated samples: Polypyrimidine Tract Binding Protein 1 (PTBP1) (Fig. 1B). We confirmed the interaction by showing higher co-immunoprecipitation of PTBP1 with RUNX1, compared to IgG, in 3 independent mouse *CM*^+^ leukemia samples (Fig. 1C). We also performed immunoprecipitations/westerns (IP/WB) using the human *CM* expressing AML cell line, ME-1, and found that RUNX1 interacts with PTBP1 in human leukemia cells, as well. To test if this interaction is unique to cells expressing the *CM* fusion gene, we performed IP/WB using a panel of human acute myeloid (AML) and acute lymphoid leukemia (ALL) cell lines. We observed co-immunoprecipitation of PTBP1 and RUNX1 in all the cell lines tested (Fig. 1D & E), indicating that the interaction between PTBP1 and RUNX1 is present in both myeloid and lymphoblastic leukemia cells.

**Figure 1.**
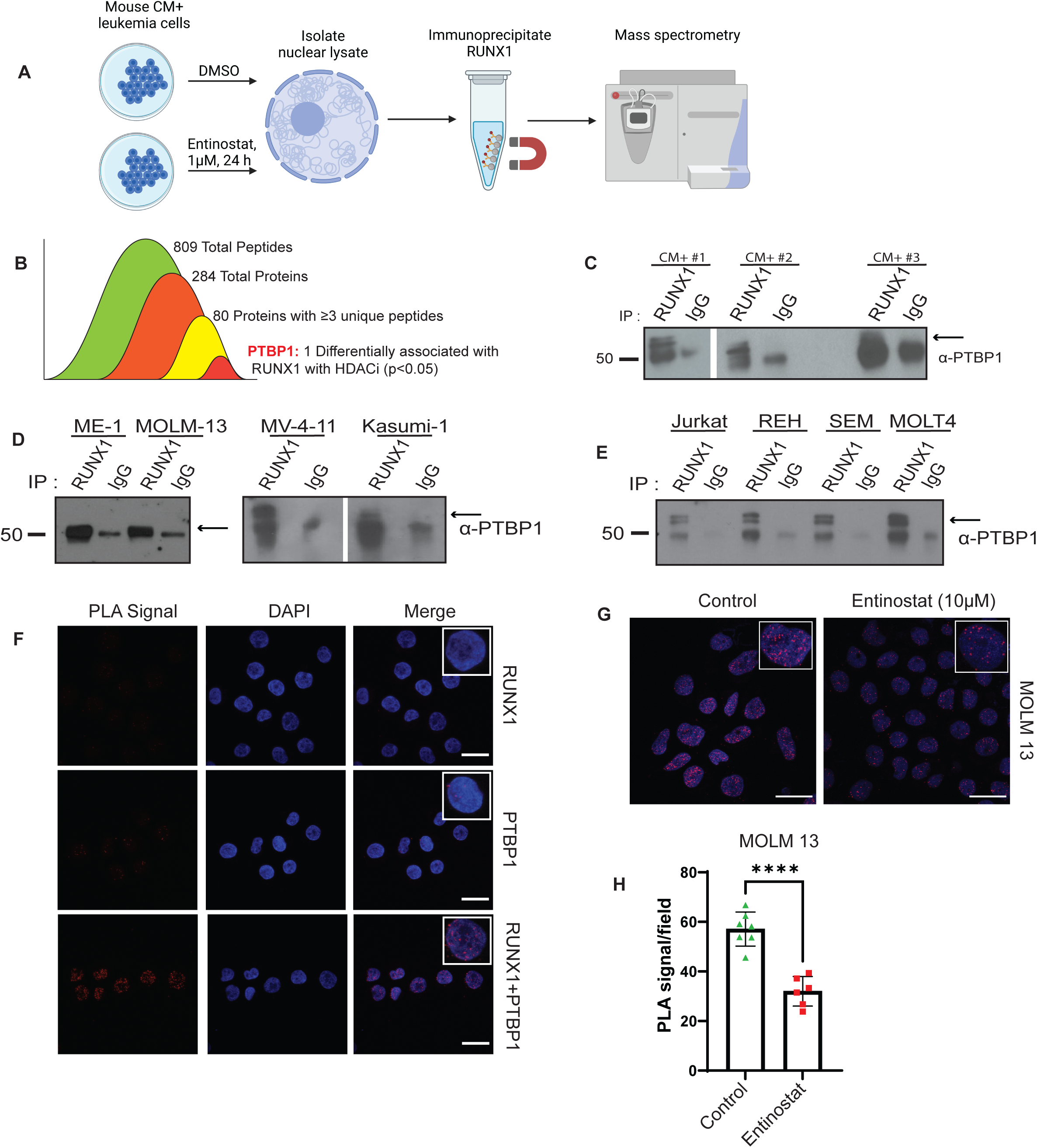
PTBP1 binds RUNX1 in a HDAC1-dependent manner: A) Schematic of experimental workflow. B) Diagram showing filtering of peptides from mass spectrometry. C) Representative western blot (WB) of lysates isolated from *CM^+^* cells, (D) human AML cell lines, or (E) human ALL cell lines immunoprecipitated with anti-RUNX1 or anti-IgG antibodies and blotted for PTBP1. F) Confocal images of Proximity Ligation Assays (PLA) performed in MOLM-13 cells using the indicated antibodies. G) Confocal images of PLAs in MOLM-13 cells treated with vehicle or 10µM entinostat for 48 hours. H) Graph quantifying PLA signal per field (minimum of 50 cells) from cells treated as in G. N>3. * = p < 0.05. **** = p < 0.0001. Scale bar = 20µm.

Both PTBP1 and RUNX1 can reside in the nucleus and the cytoplasm. To determine in which cellular compartment the proteins are interacting, we performed proximity ligation assays (PLA). We observe very few PLA puncta in RUNX1 or PTBP1 only antibody controls but observed multiple puncta in the nuclei of cells with both PTBP1 and RUNX1 antibodies (Fig. 1F). To confirm that HDAC1 activity regulates the interaction between RUNX1 and PTBP1, we performed PLA on MOLM-13 and REH cells treated for 48 hours with vehicle or entinostat. We found that in both cell lines (Fig. 1G & 1H, Supplementary Fig. 1A & B) there were significantly fewer PLA puncta in cells treated with entinostat, consistent with our mass spectrometry results. These findings indicate that RUNX1 and PTBP1 interact in the nucleus of both AML and ALL cells, and this interaction requires HDAC1 activity.

### The N-terminus of PTBP1 and the C-terminus of RUNX1 are required to interact

To ascertain protein domains responsible for the interaction between RUNX1 and PTBP1, we used expression constructs with full length, wild type (WT) FLAG tagged RUNX1, and a series of point and deletion mutations found in patients with Familial Platelet Disorder with Associated Myeloid Malignancy (FPDMM) (Fig. 2A) (28–30, 42). We observed co-IP of Myc-PTBP1 with WT Flag-RUNX1, Flag-RUNX1-G365R and Flag-RUNX1-Y407X, but not with Flag-RUNX1-R232X, Flag-RUNX1-Y287X, and RUNX1-R320X (Fig 2B & C). This implies that PTBP1binds between amino acids 320 and 407 of RUNX1. To test which region of PTBP1 is required to bind RUNX1, we generated a PTBP1 truncation mutant that contains only the N-terminal region and the first RNA recognition motif (RRM), Myc-PTBP1-RRM1 (Fig. 2A). We observed co-IP of WT Flag-RUNX1 with full length Myc-PTBP1 and Myc-PTBP1-RRM1, implying that the N-terminus of PTBP1 binds RUNX1 (Fig. 2D). Collectively, these results indicate that the N-terminus of PTBP1 and the C-terminus of RUNX1 are required for the interaction between RUNX1 and PTBP1.

**Figure 2.**
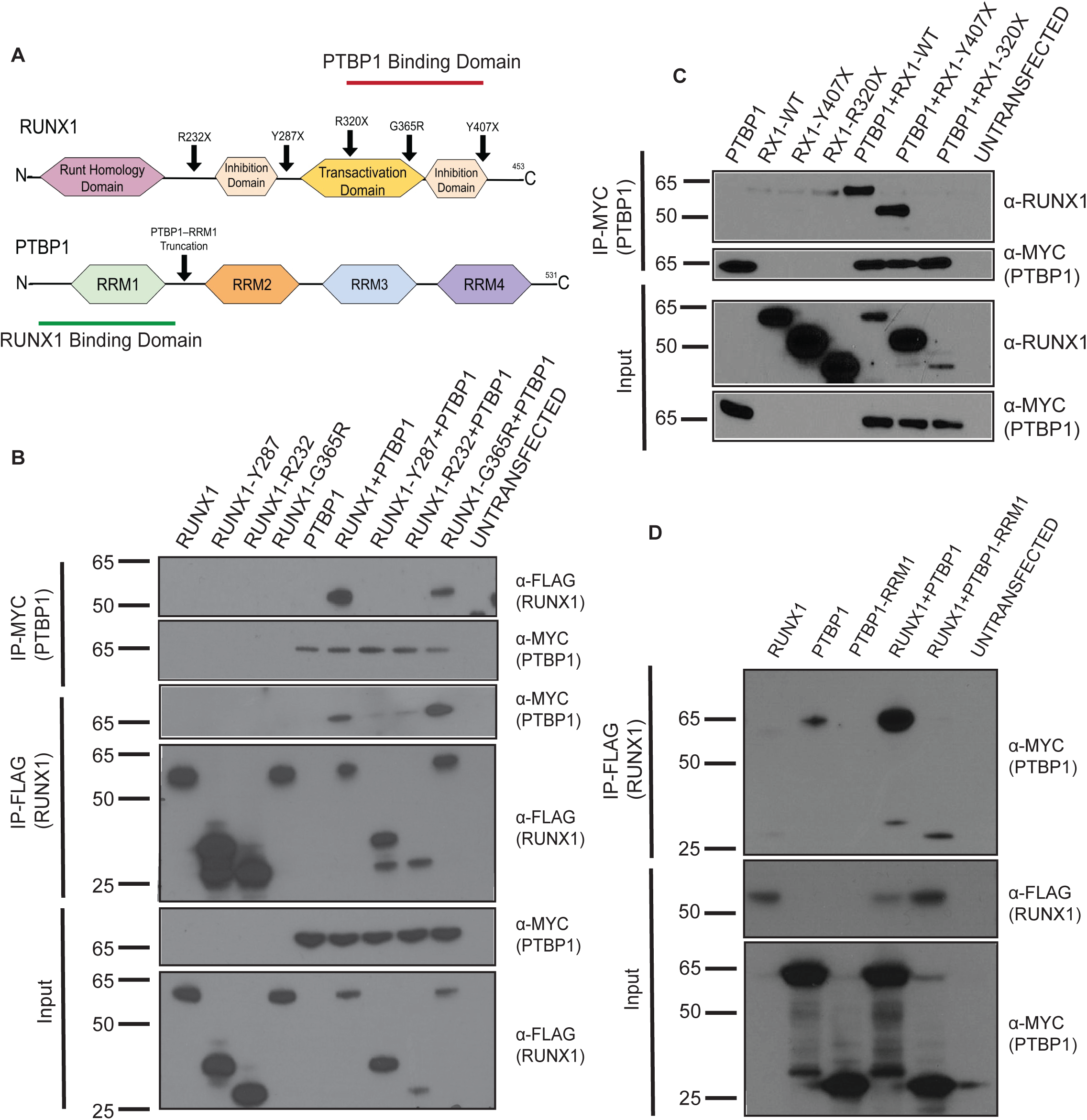
The N-terminus of PTBP1 binds the C-terminus of RUNX1: A) Schematic representation of RUNX1 and PTBP1 proteins showing binding domains, and the relevant mutations indicated with arrows. B-D) Representative western blots from HEK293T cells transfected with expression constructs for the indicated proteins, immunoprecipitated (IP’d) and probed with the indicated antibodies. Each experiment was performed at least 3 times.

### The PTBP1-RUNX1 interaction occurs in healthy hematopoietic stem and early progenitor cells

RUNX1 is known to have roles in both leukemia and healthy hematopoietic stem and progenitor cells (HSPCs). To address whether RUNX1 binds PTBP1 in these cells, we compared the expression of PTBP1 in leukemia cells and HSPCs. Using publicly available gene expression data, we found that *PTBP1* is expressed at similar levels in leukemia cells and healthy HSPCs but decreases with differentiation (Fig.3A). To examine PTBP1 protein expression, we performed WBs with bone marrow (HBM) and CD34+ cells from healthy volunteers, and primary AML patient samples. We found that PTBP1 is undetectable in HBM but highly expressed in CD34+ cells. PTBP1 was detectable in most patient samples, but its expression levels showed considerable variation (Fig. 3B). To test if RUNX1 binds PTBP1 in healthy hematopoietic cells, we performed PLA on mouse lineage depleted (lin-) bone marrow cells and human CD34+ cells. Primary mouse *CM*^+^ cells and patient AML cells were included for comparison. We observed significantly more PLA puncta in *CM^+^* leukemia compared to wild type lin-bone marrow cells (Fig. 3C & D). Human CD34+ as well as primary AML cells showed substantial levels of PLA puncta (Fig. 3E).

**Figure 3.**
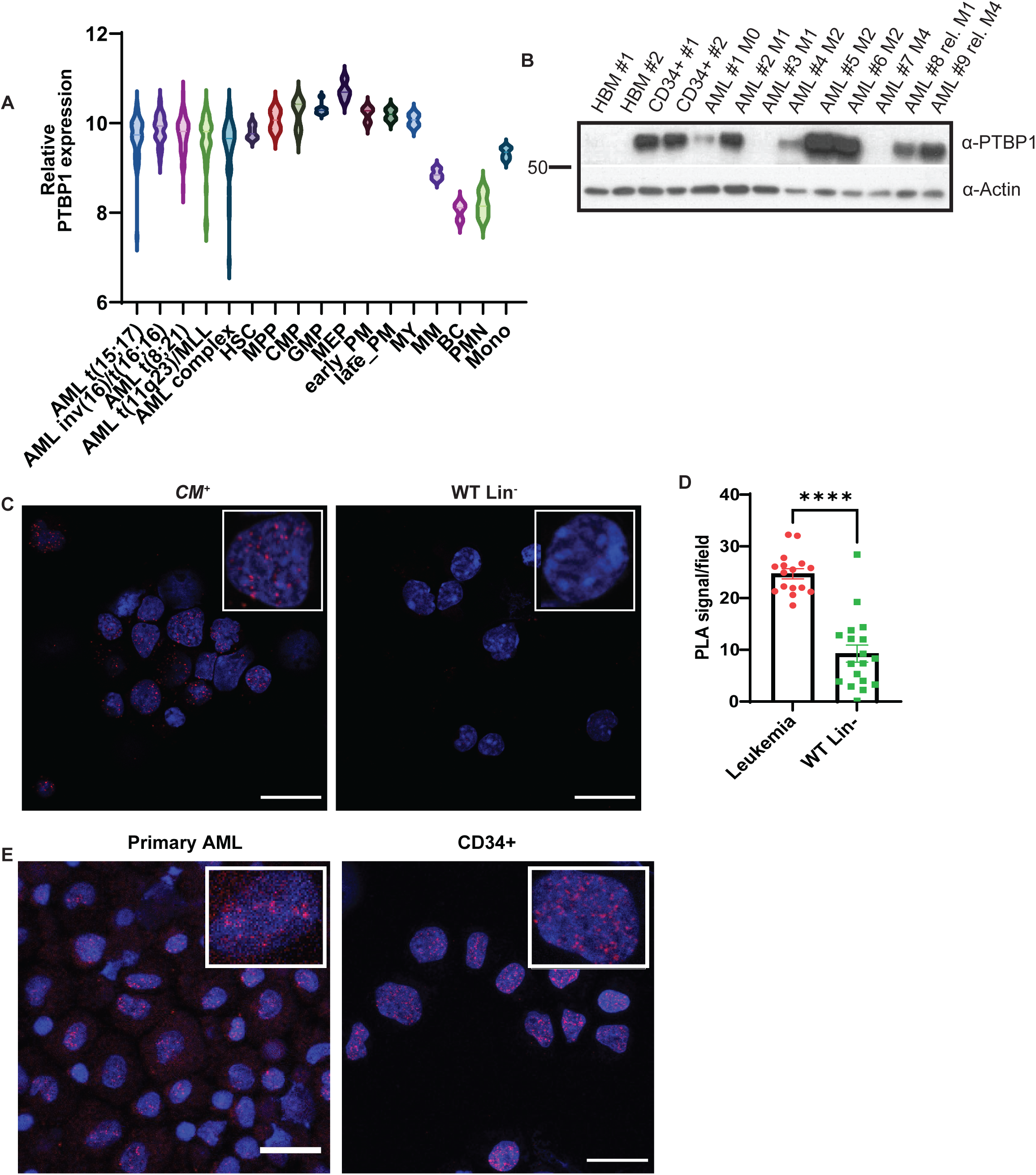
PTBP1-RUNX1 interaction occurs in healthy hematopoietic stem and early progenitor cells: A) Violin plot depicting relative *PTBP1* expression in human hematopoietic populations and AML subtypes (HSC-Hematopoietic Stem Cell; MPP-Multipotent Progenitor; CMP- Common Myeloid Progenitor; GMP-Granulocyte-Monocyte Progenitor; MEP-Megakaryocyte-Erythrocyte Progenitor; early PM-Early Promyelocyte; late PM-Late Promyelocyte; MY-Myelocyte; MM-Metamyelocytes; BC-Band cell; PMN-Polymorphonuclear cells; Mono-Monocytes). B) Western blot (WB) of PTBP1 protein in healthy bone marrow (HBM) and sorted CD34^+^ cells from healthy individuals, and primary diagnosis and relapse (rel) AML samples of the indicated FAB stage. C) Representative confocal images and D) graph of PLA intensity/field (minimum of 50 cells) in mouse CM^+^ leukemia cells and WT lineage depleted (Lin-) cells. E) Representative confocal images of PLA performed with RUNX1 and PTBP1 antibodies in healthy human CD34+ cells and the human AML patient sample 6406. N=3. **** = p < 0.0001. Scale bar - 20µm.

As our expression data and work in genetic mouse models indicate a role for PTBP1 in healthy hematopoietic stem cells, we asked whether PTBP1 expression promotes leukemia stem cell (LSC) activity (16). For this, we used mouse *CM^+^*leukemia cells expressing GFP from the Rosa26 locus (*CM^+^, GFP^+^*), and took advantage of the natural variability in PTBP1 expression levels in primary leukemia cells (Supplementary Fig. 2A). *CM^+^, GFP^+^*leukemia cells were sorted for GFP, and WBs were used to identify mouse leukemia samples with PTBP1 expression above (PTBP1 high) and below (PTBP1 low) the median expression level (Supplementary Fig. 2B). Equal numbers of cells from 3 independent PTBP1 high and PTBP1 low samples were plated for colony formation assay. We observe significantly more colonies in samples expressing high levels of PTBP1, implying that increased PTBP1 expression is associated with increased LSC activity (Supplementary Fig. 2C & D).

### PTBP1 co-localizes with RUNX1 at promoter regions

To determine if PTBP1 co-localizes with RUNX1 on chromatin, we performed CUT&Tag (Cleavage Under Targets and Tagmentation) using antibodies against, RUNX1, PTPB1, and the appropriate IgG controls. We observed extensive overlap between PTBP1 and RUNX1 peaks, withRUNX1 signal at nearly all PTBP1 peaks and vice-versa (Fig. 4A). We categorized peaks into three groups: the largest category being those with equivalent binding of both PTBP1 and RUNX1, with fewer regions showing either stronger PTBP1 (PTBP1>RUNX1) or stronger RUNX1 (PTBP1<RUNX1). To identify the types of genomic regions bound by PTBP1 and RUNX1, we compared enrichment of signal in peaks at annotated Transcription Start Site (TSS), genes, and intergenic regions. We found that for all three categories of peaks, PBTP1 and RUNX1, were significantly enriched at TSS and gene bodies compared to Monte Carlo permutations (Fig. 4B,C). To test whether PTBP1 and RUNX1 binding is associated with active transcription, we compared our CUT&Tag data to publicly available RNA Polymerase II and histone modification chromatin occupancy data from MOLM-13 cells (43). Regions bound by both PTBP1 and RUNX1 show strong enrichment of RNA Pol II and histone modifications associated with active transcription (H3K9ac, K3K27ac, H3K4me1, and H3K4me3), but not marks of transcriptional repression (H3K27me3) (Fig. 4D, E, & Supplementary Fig. 3A-C). Further characterization of chromatin states using ChromHMM (44), revealed that PTBP1 and RUNX1 together are significantly enriched at strongly transcribing promoters (Fig. 4F, G, Supplementary Fig. 3D).

**Figure 4.**
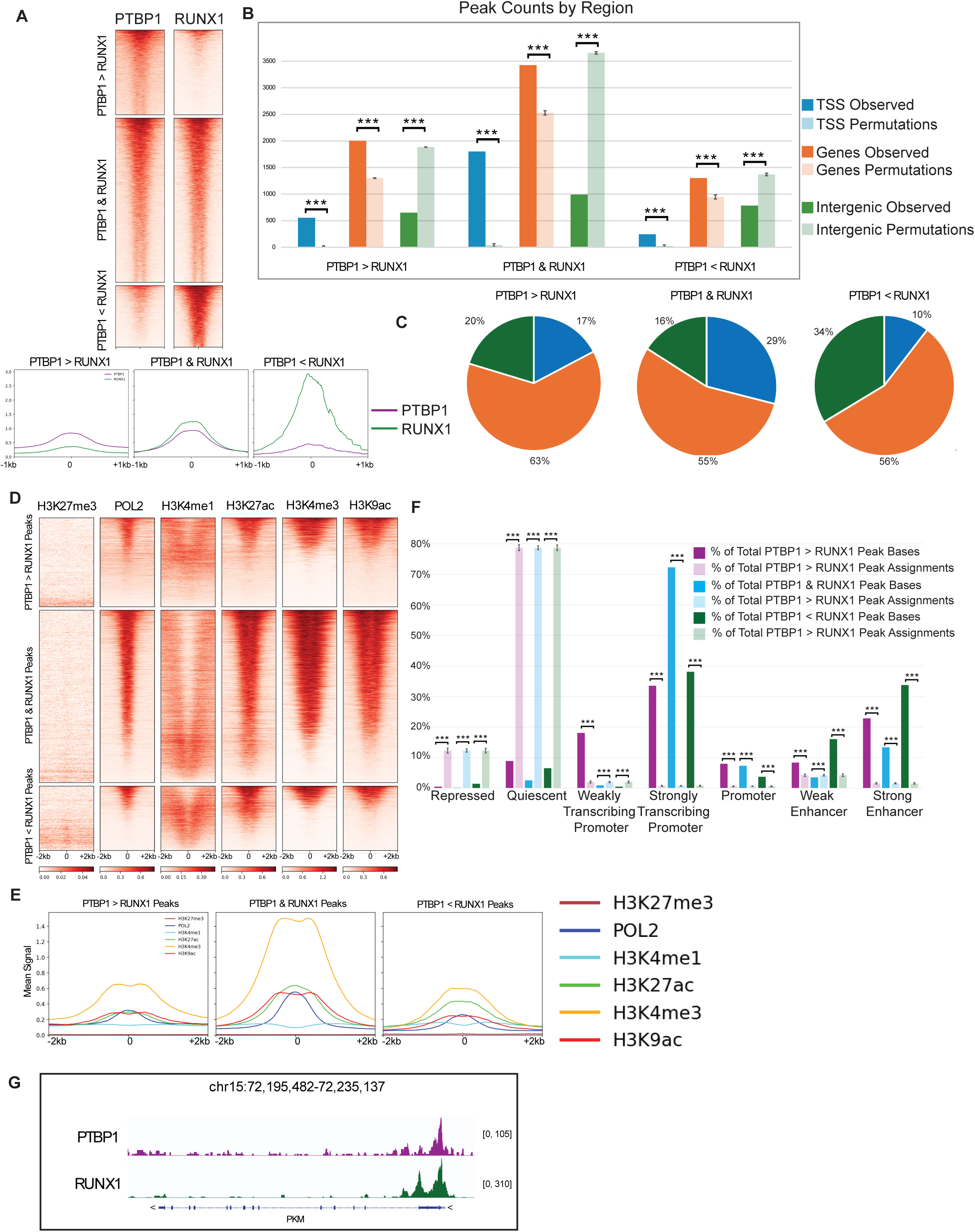
PTBP1 co-localizes with RUNX1 at promoter regions: A) Heatmaps and mean signal intensities from PTBP1 and RUNX1 CUT&Tag, centered on the peak, with 0 indicating the midpoint (±1kb) grouped into regions with PTBP1 high (PTPB1>RUNX1), PTBP1 and RUNX1 high (PTBP1 & RUNX1), and RUNX1 high (PTBP1<RUNX1). Peaks were called using MACS3 and the three groups of bound regions defined using MAnorm. Heatmaps were sorted by decreasing signal intensity. B) Location of peaks relative to annotated genomic loci; lightly colored bars display results of Monte Carlo permutations (n=1000), demonstrating likelihood against random relocation of peaks. TSS-Transcription Start Site. C) Pie chart representing occupancy of either RUNX1, or PTBP1, or both at annotated genomic loci. D) Heatmap and E) mean signal intensities of RNA Polymerase 2 (POL2) and histone modifications, specifically H3K27me3, H3K4me1, H3K27ac, H3K4me3, H3K9ac centered on the midpoint of the peak (±2kb). F) Percent distribution of peaks over ChromHMM-generated states; lightly colored bars display results of Monte Carlo permutations (n=1000), demonstrating likelihood against random relocation of peaks. G) IGV snapshot showing high PTBP1 & high RUNX1 signal at PKM gene TSS. *** = p≤ 0.001.

### Loss of PTBP1 induces widespread changes in splicing and gene expression

The co-localization of PTBP1 with RUNX1 at active promoters raises the possibility that PTBP1 has a role in the expression and splicing of target genes. To test this possibility, we used a doxycycline (dox) inducible shRNA against PTBP1 (sh*PTBP1*) or a scrambled shRNA (sh*SCR*) (45). These lentiviral constructs constitutively express mCherry and express ΔLNGFR along with the shRNA after dox treatment. After confirming efficient KD of PTBP1 (Supplementary Fig. 4), we performed long-read RNA sequencing in control and PTBP1 KD MOLM-13 cells. Principal component analysis showed that the sh*PTBP1* samples cluster together, distinct from the control sh*SCR* samples (Fig. 5A). We also found that loss of PTBP1 induced changes in splicing, with the most common changes involving the use of alternative transcriptional start sites (Alt TSS) (25%), alternative termination sites (Alt TTS) (23%), and skipped exons (24%) (Fig. 5B), with 131 significant differential isoforms associated with 83 genes (Fig. 5C). Analysis of our CUT&Tag data for the isoform switched genes shows this gene set is enriched for RUNX1, PTBP1, and RNA Pol II binding, as well as histone modifications associated with active transcription, at the TSS (Supplementary Fig. 5A & B). Comparing the list of isoform switched genes to publicly available PTBP1 eCLIP data, we found that the majority of these transcripts (77 of 83) are bound by PTBP1, indicating they are PTBP1 splicing targets (Supplementary Fig. 5C-E) (46).

**Figure 5.**
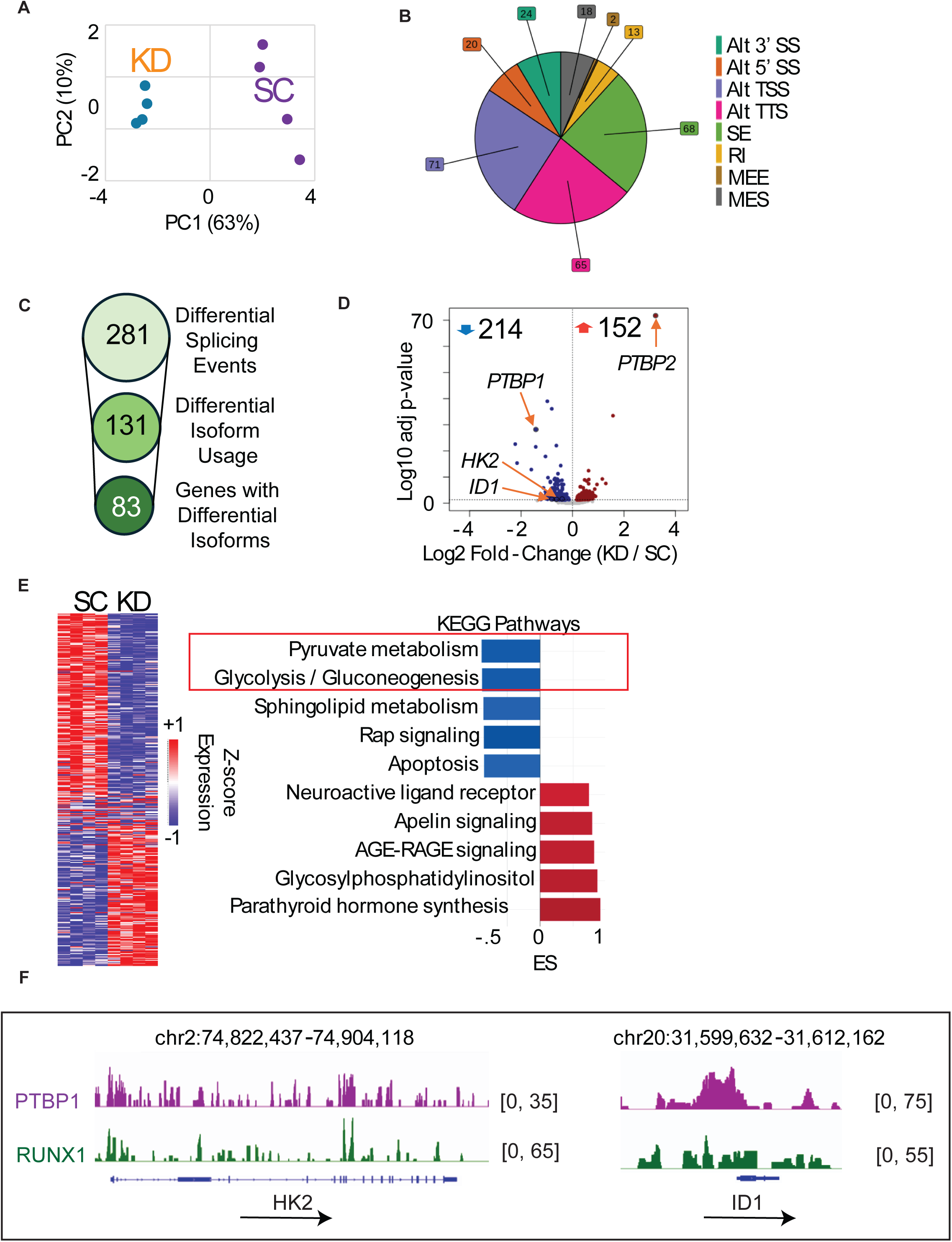
Loss of PTBP1 induces widespread changes in splicing and gene expression: A) Principal component analysis of MOLM-13 sh*SCR* (SC) and sh*PTBP1* (KD) cells from long-read RNA sequencing data. B) Pie chart revealing altered splicing events in response to PTBP1 KD in MOLM-13s (Alt 3’ SS – Alternative 3’ Splice Site; Alt 5’ SS – Alternate 5’ Splice Site; Alt TSS – Alternate Transcription Start Site; Alt TTS – Alternate Transcription Termination Site; SE – Skipped Exon; RI – Retained Intron; MEE – Mutually Exclusive Exon; MES – Multiple Exon Skipping). C) Analysis of the splicing events in (B.) revealing genes with differential isoform expression in response to PTBP1 KD in MOLM-13s. D) Volcano plot showing differentially expressed genes in response to PTBP1 KD in MOLM-13s. Highlighted genes include Inhibitor of Differentiation 1 (*ID1*), Hexokinase-2 (*HK2*), *PTBP1*, and *PTBP2*. E) Analysis of differentially expressed genes by EnrichR showing top KEGG pathways affected after PTBP1 KD. F) Integrated Genomics Viewer (IGV) tracks depicting the binding of RUNX1 and PTBP1 at *HK2* and *ID1* gene loci. N≥3.

In addition to altered isoform usage, PTBP1 KD resulted in the downregulation of 214 and upregulation of 152 genes, with the most highly upregulated gene being the fellow PTBP family member, PTBP2. Among the downregulated genes, we identified of key metabolic genes including Hexokinase-2 (*HK2*), Inhibitor of DNA Binding 1 (also known as Inhibitor of Differentiation 1) (*ID1*) and Solute Carrier 2A1 (*SLC2A1*), as well as PTBP1. (Fig. 5D). Functional categorization and overrepresentation analysis of Kegg pathways revealed that the genes downregulated with PTBP1 KD are associated with metabolic pathways critical for leukemia cells growth, including pyruvate metabolism and glycolysis (Fig. 5E). Our CUT&Tag data identified RUNX1 and PTBP1 co-localization at several deregulated genes associated with these pathways, including *ID1*, *HK2* (Fig. 5F), and *SLC2A1*, which encodes Glucose Transporter 1 (GLUT1) (Supplementary Fig. 6B).

### PTBP1 promotes leukemia cell survival by regulating expression of metabolic genes

Downregulation of genes associated with glycolysis and pyruvate metabolism with PTPB1 KD implies a role for PTBP1 in regulating leukemia cell growth. To test this, we treated sh*PTBP1* and control sh*SCR* cells with dox for 2 weeks and performed growth assays. We observed a significant reduction in cell numbers in *PTBP1* KD cells compared to control cells (Fig. 6A & B). To test if the decreased growth is associated with differentiation to a post-mitotic, mature myeloid phenotype, we examined the expression CD11b and CD15. We found that 9 days of dox treatment induced a statistically significant increase in CD15 expression in sh*PTBP1* cells, as compared to sh*SCR* cells (Fig. 6C & D). To test if cell death also contributes to the growth defect in PTBP1 KD cells, we used flow cytometry. At day 7 of PTBP1 KD, we did not observe any effect on viability (Fig. 6E). In contrast, at day 21 there was significantly increased Annexin V+ staining in PTBP1 KD cells indicating that prolonged knockdown of PTBP1 induces apoptosis (Fig. 6F & G).

**Figure 6.**
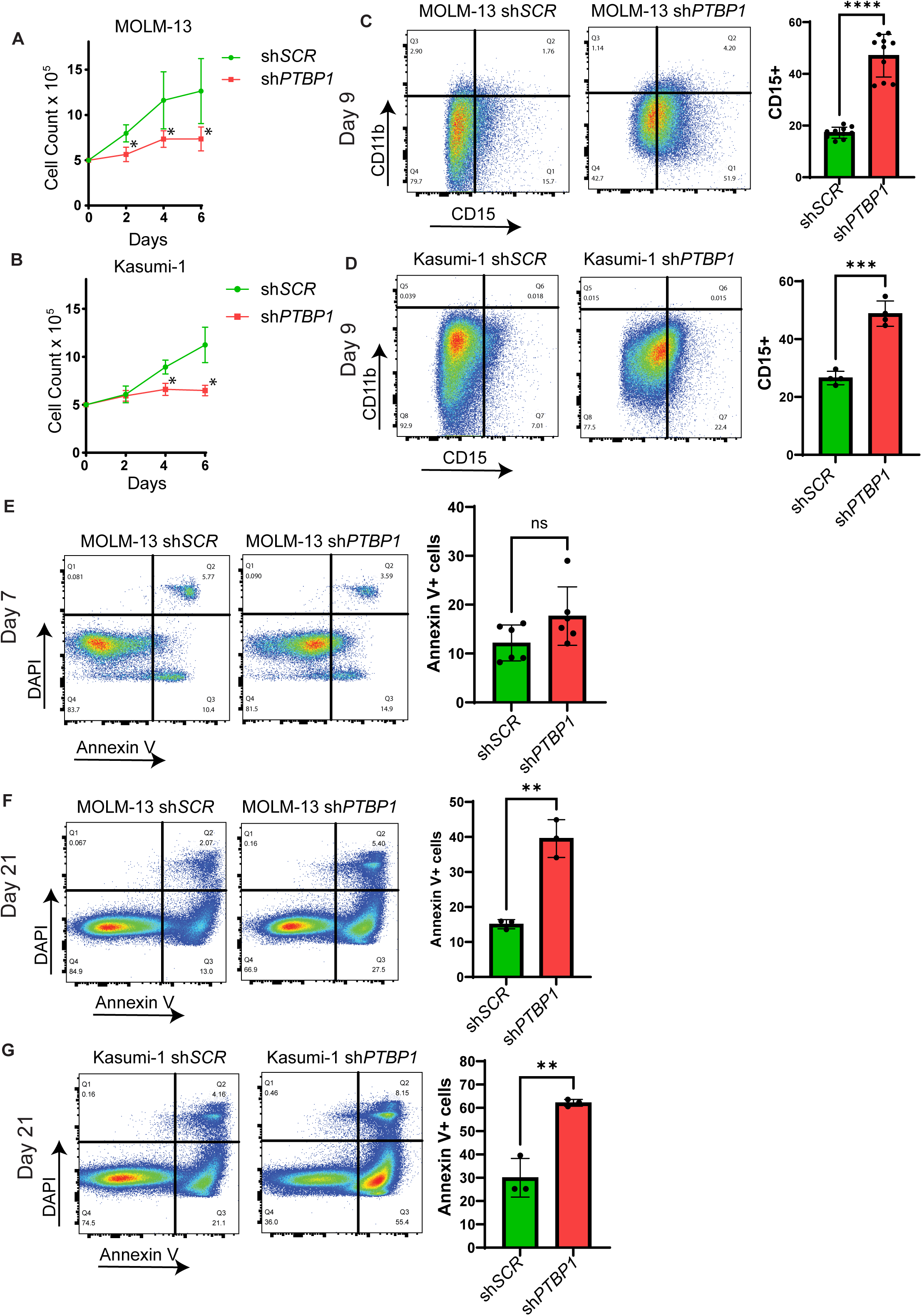
PTBP1 promotes leukemia cell survival by regulating expression of metabolic genes: A) Growth curve of MOLM-13 sh*SCR* and sh*PTBP1* cells. B) Growth curve of Kasumi-1 shS*CR* and sh*PTBP1* cells. C) Representative plots and quantification from flow cytometry analysis of CD15 and CD11b of MOLM-13 sh*SCR* and sh*PTBP1* cells and D) Kasumi-1 sh*SCR* and sh*PTBP1* cells. E) Representative plots and quantification of Annexin V+ cells in MOLM-13 sh*SCR* and sh*PTBP1* cells on day 7 of doxycycline induction. F) Representative plots and quantification of flow cytometry analysis of Annexin V and DAPI on day 21 of doxycycline induction in MOLM-13 sh*SCR* and sh*PTBP1* cells and G) Kasumi-1 sh*SCR* and sh*PTBP1* cells. N≥3. * = p < 0.05, ** = p < 0.01. **** = p < 0.0001.

To determine if growth defects caused by PTBP1 KD are accompanied by metabolic dysfunction, we first performed western blot analysis of metabolism related proteins found deregulated in our RNA-seq analysis. We found that ID1 and HK2 showed significantly decreased expression after 7 days of KD (Fig. 7A & B). At this time point, PTBP1 KD cells showed significantly lower lactate release into the media, which is consistent with decreased glycolysis (Fig. 7C). At day 14 of PTBP1 KD, we also observed a trend towards decreased GLUT1 expression. Using a flow cytometry-based glucose uptake assay, we observed a significant reduction in glucose uptake specifically in viable, DAPI-, sh*PTBP1* cellsat this time point(Fig. 7D & E). Collectively, these results indicate that PTBP1 regulates the expression of key metabolic genes required for efficient glycolysis and survival in leukemia cells.

**Figure 7.**
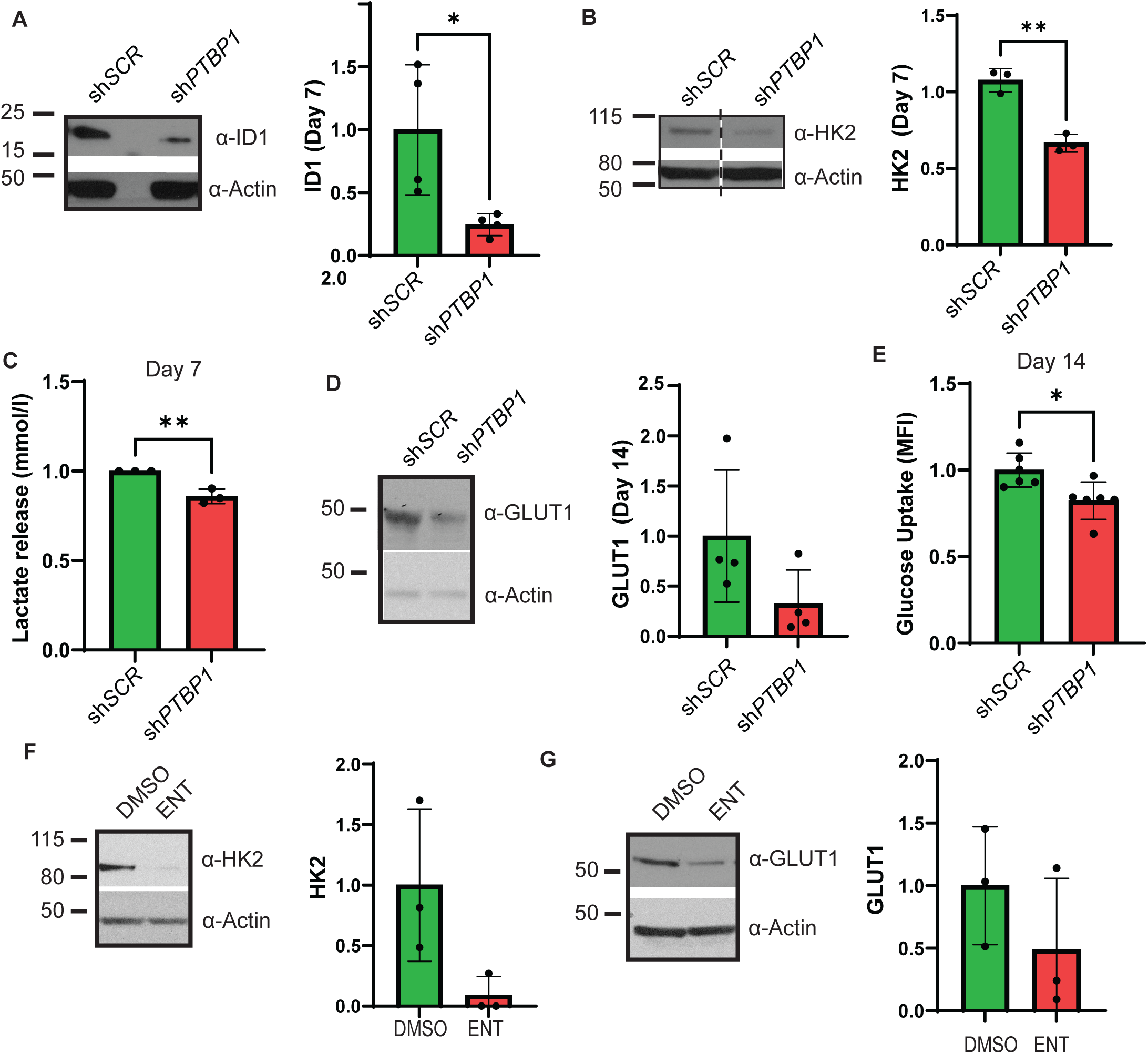
Loss of PTBP1 causes metabolic defects in leukemia cells: A) Representative WB and quantification of relative abundance of Inhibitor of DNA binding 1 (ID1) at day 7 of PTBP1 KD in MOLM-13s. B) Representative WB and quantification of relative abundance of Hexokinase-2 (HK2) at day 7 of PTBP1 KD in MOLM-13s. Dashed line indicates non-adjacent lanes from the same gel. C) Quantification of lactate from lactate release assays in MOLM-13 sh*SCR* and sh*PTBP1* cells at day 7 of KD. D) Representative WB and quantification of relative abundance of Glucose transporter-1 (GLUT1) on day 14 of PTBP1 KD in MOLM-13s. E) Quantification of mean fluorescence intensity (MFI) from glucose uptake assays on day 14 of doxycycline treatment in MOLM-13 sh*SCR* and sh*PTBP1* cells. F) Representative WB and quantification of relative abundance of HK2 And G) GLUT1 in MOLM 13 cells treated with 5 µM Entinostat (ENT) or DMSO for 48 hours. N≥3. * p ≤ 0.05. ** = p < 0.01.

To begin to address whether the interaction with RUNX1 may contribute to PTBP1’s role in the expression of these genes, we treated MOLM-13 cells with 5µM entinostat for 48 hours to disrupt the RUNX1/PTBP1 interaction. We observed a trend towards decreased HK2 and GLUT1 in entinostat treated cells, consistent with a role for the RUNX1/PTBP1 interaction in promoting expression of their target genes (Fig. 7F & G).

### PTBP1 KD induces DNA damage and sensitizes leukemia cells to chemotherapy

Deletion of the RUNX1 C-terminus, the region that interacts with PTBP1, has been shown to induce DNA damage (47). To determine whether KD of PTBP1 has a similar effect in leukemia cells, we examined expression of the DNA damage marker, phosphorylated γH2A.X, in PTBP1 KD and control cells. We observed elevated levels of phospho-γH2A.X in PTBP1 KD cells (Fig. 8A & B). To test if the increased DNA damage sensitizes PTBP1 KD cells to chemotherapy, we treated control and PTBP1 KD cells with 5µM of cytarabine for 3 days. We observed a significant increase in the Annexin V+ population in PTBP1 KD cells as compared to control, indicating that loss of PTBP1 sensitizes leukemia cells to chemotherapy (Fig. 8C & D).

**Figure 8.**
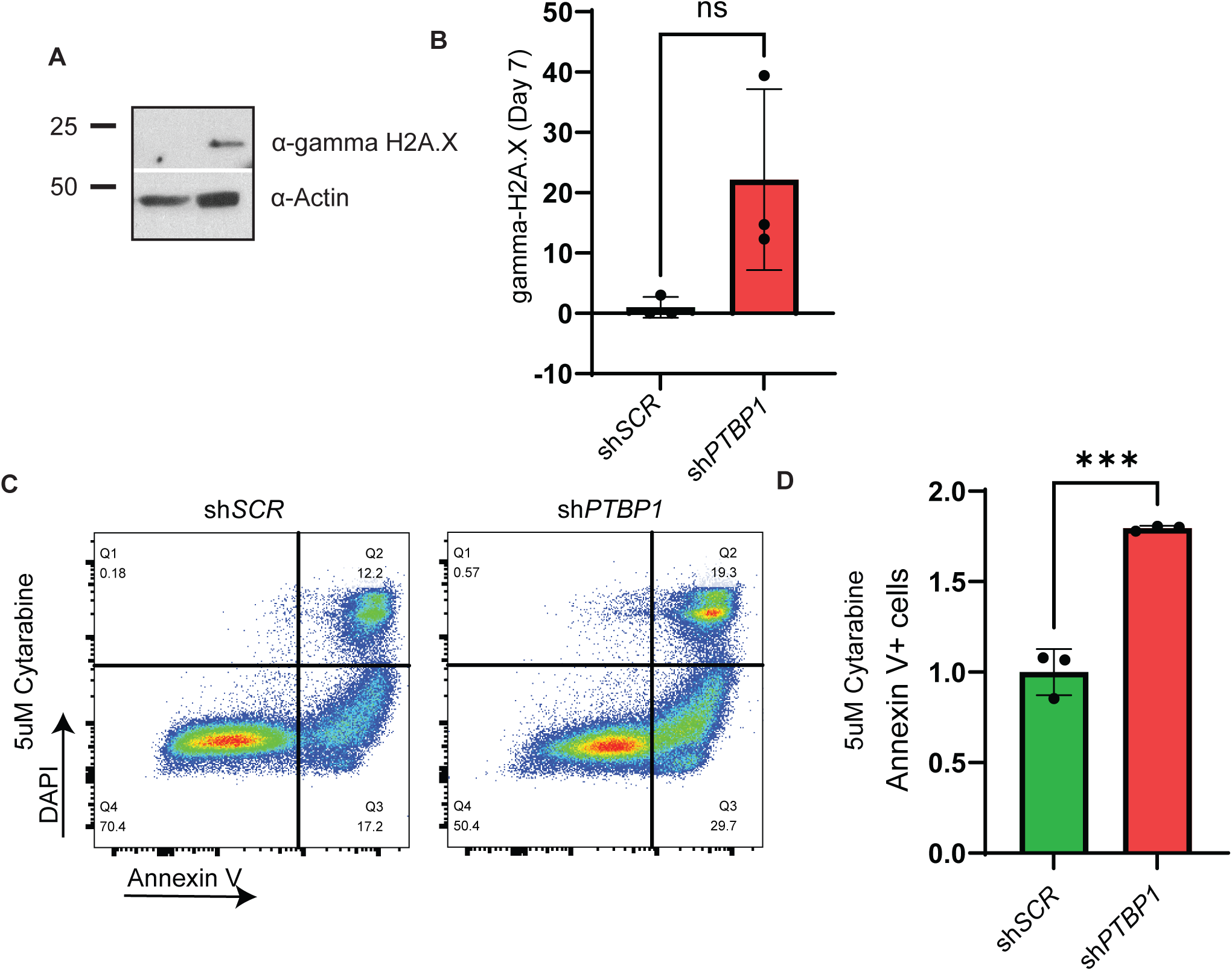
PTBP1 KD sensitizes leukemia cells to chemotherapy: A) Representative WB and B) quantification of (S139) phosphorylated ɣH2A.X (S139) levels after 7 days of PTBP1 KD in MOLM-13 cells. C) Representative plots of flow cytometry analysis and D) quantification of Annexin V and DAPI staining in MOLM-13 sh*SCR* and sh*PTBP1* cells treated with 5µM cytarabine for 72 hours after 7 days of PTBP1 KD. N≥3. *** = p < 0.001.

Collectively, our findings indicate that RUNX1 and PTBP1 interact in an HDAC1 dependent manner at the promoters of key metabolic genes critical for leukemia cells survival. This work supports a model in which recruitment of PTBP1 to RUNX1 target genes promotes their efficient expression and maintains robust glycolytic output, highlighting a previously unrecognized aspect of RUNX1’s central role in leukemia pathogenesis.

## Discussion

HDAC1 is canonically considered a repressor of gene expression but can also mediate protein interactions. Previous work from our lab showed that HDAC1 is required for active transcription of a subset of RUNX1 target genes in leukemia cells. To address this contradiction, we tested whether inhibition of HDAC1 alters the RUNX1 interactome. We found that PTBP1 binds RUNX1 in an HDAC1-dependent manner and is required for expression of genes bound by both factors, providing a potential mechanism for HDAC1’s non-histone role in gene regulation. In addition, we identified the regions in RUNX1 and PTBP1 required for this interaction. Interestingly, both domains include multiple lysine residues that could be targets of HDAC1 mediated deacetylation. In the case of PTBP1, lysines (K92 & K94) in the RUNX1 interaction domain have been shown to be acetylated, providing further support for this potential mechanism (48).

We also showed that both RUNX1 and PTBP1 exhibit a strong bias for binding the transcription start site of actively transcribed genes, including those involved in metabolism. Previous work has implicated PTBP1 in the regulation of glycolysis by altering the splicing of pyruvate kinase (PKM) in leukemia (18) (22). In the current study, we identify additional metabolic genes dependent on PTBP1 for expression, including *SLC2A1*, which encodes the glucose transporter GLUT-1, Hexokinase 2 (*HK2*), an enzyme in the glycolysis pathway, and Inhibitor of DNA Binding 1 (*ID1),* an upstream regulator of glycolysis (49). We found that loss of PTBP1 induced decreased expression of these factors at the RNA and protein level and caused metabolic defects. It is notable that we observed decreased expression of these genes, rather than the presence of alternative or mis-spliced isoforms. As PTBP1 is not thought to have any transactivation activity on its own, the most likely explanation for this observation is that these transcripts were subject to nonsense mediated decay. Another explanation is that PTBP1 promotes the stability of these transcripts, a PTBP1 function observed in other cancers (50). Importantly, the changes in expression and the associated metabolic defects were observed prior to induction of apoptosis, implying that the decreased expression of the RUNX1/PTBP1 target genes is the cause, and not the consequence, of cell death in PTBP1 KD cells.

Our study demonstrates a novel facet of RUNX1’s role in leukemia through interaction with PTBP1 and the regulation of metabolism. Our findings lead us to propose the following mechanism: PTBP1, which does not have a DNA binding domain, binds RUNX1 and is recruited to the promoter regions to drive expression of genes essential for leukemia cell survival. As a result, loss of PTBP1 leads to decreased expression of these target genes, leading to reduced metabolic output and ultimately cell death (See Graphical Abstract). Collectively, our results provide new insights into the intricacies of co-transcriptional splicing in leukemia and hematopoiesis, further underscoring the central role RUNX1 plays in both these processes.

## ACKNOWLEDGEMENTS

This work was supported by R01 CA244900 to R.K.H., R35 GM147467 to M.J.R, R00GM138920 to S.B, UNMC graduate assistantship/fellowships to A.D. and C.L and T32 CA009476 to C.L. The UNMC Flow Cytometry Research Facility, Microscopy Core Facility and Genomic Core Facility are supported by the Nebraska Research Initiative (NRI), P30 CA036727, P20 GM103427, P30 GM106397, P20GM130447, S10RR02730, S10OD030486, and P20 GM121316.

## AUTHOR CONTRIBUTIONS

A.D. and R.K.H. conceived of and designed the study. A.E., S.B., and M.J.R helped design and analyze the long-read transcriptomic and CUT&Tag experiments. A.D., C.L., R.W., K.T.R., J.P, S.P., K.D. and S.S. performed and analyzed experiments. A.D., A.E., R.W., S.B., M.J.R, and R.K.H. wrote the manuscript.

## COMPETING INTERESTS

The authors have no competing interests to declare.

## DATA AVAILABILITY STATEMENT

All proteomic and genomic data generated in this study will be deposited in the appropriate publicly accessible databases by the time of publication.

## Notes

**Competing Interests:** This work was supported by R01 CA244900 to R.K.H., R35 GM147467 to M.J.R, R00GM138920 to S.B, UNMC graduate assistantship/fellowships to A.D. and C.L and T32 CA009476 to C.L. The UNMC Flow Cytometry Research Facility, Microscopy Core Facility and Genomic Core Facility are supported by the Nebraska Research Initiative (NRI), P30 CA036727, P20 GM103427, P30 GM106397, P20GM130447, S10RR02730, S10OD030486, and P20 GM121316.

### Competing Interest Statement

The authors have declared no competing interest.

