## Supplemental Files for "The Splicing Factor PTBP1 interacts with RUNX1 and is Required for Leukemia Cell Survival"

### Supplementary Figure Legends

**Supplementary Figure 1**. A) Confocal images of PLAs in REH cells treated with vehicle or 5μM entinostat for 48 hours. B) Graph quantifying PLA signal per field (minimum of 50 cells). N>3. **** = $p < 0.0001$. Scale bar - 20μm.

**Supplementary Figure 2.** A) Representative WB and B) graph of relative PTBP1 levels standardized to actin from 7 independent $CM^+$ mouse leukemia cells. C) Representative images (10x magnification) and D) graph of colony numbers from colony assays of PTBP1 high and low $CM^+$ mouse leukemia cells from (A).

**Supplementary figure 3.** A) Heatmap and B) profile of all ChIP-seq, CUT&RUN, and CUT&Tag signals over Transcription Start Sites (TSSs) (± 2kb) noted as overlapping with any PTBP1 or RUNX1 peaks. C) Spearman correlation plot of signal over TSS overlapping with PTBP1 or RUNX1 peaks +/-500bp. D) CUT&Tag (PTBP1, RUNX1), CUT&RUN (H3K27me3), and ChIP-seq (Pol2, H3K4me1, H3K27ac, H3K4me3, H3K9ac) signals at Pyruvate Kinase M1/2 (PKM) gene locus.

**Supplementary figure 4.** A) Schematic depicting the *PTBP1* shRNA construct. B) Representative WB and quantification of PTBP1 levels normalized to actin in 3 human AML cell lines transduced with the sh*SCR* and sh*PTBP1*.

**Supplementary figure 5.** A) Heatmap and B) profile showing binding of PTBP1, RUNX1, RNA Pol II and histone modifications at Transcription Start Sites (TSSs) of 83 isoform switched genes. C) Figure displaying counts of 83 isoform switched genes that overlap with PTBP1 eClip peaks. D) PTBP1 eCLIP peak signal and E) Quantification of eCLIP peaks over 83 isoform switched genes compared to 83 randomly selected genes.

**Supplementary figure 6.** A) Integrated Genomics Viewer (IGV) tracks demonstrating CUT&Tag (PTBP1, RUNX1), CUT&RUN (H3K27me3), and ChIP-seq (Pol2, H3K4me1, H3K27ac, H3K4me3, H3K9ac) signals over Hexokinase-2 (HK2) locus. B) IGV tracks showing CUT&Tag (PTBP1 and RUNX1) signal at Glucose transporter-1 (SLC2A1) locus.

Supplementary fig 1

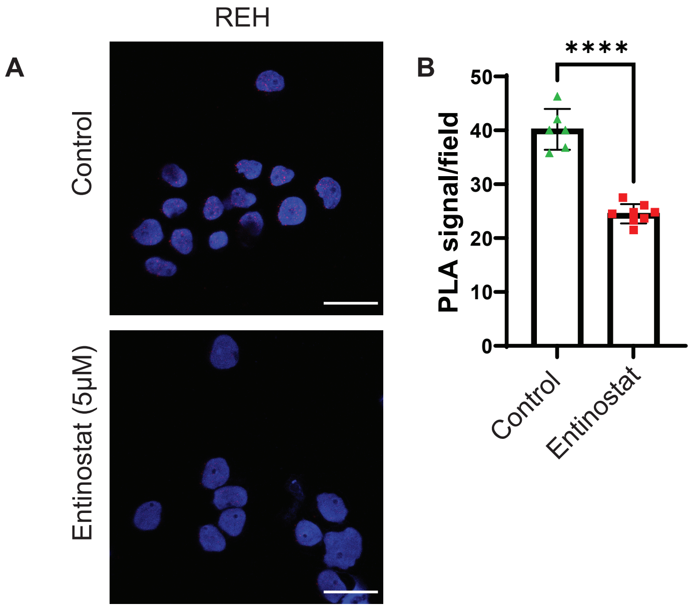

Supplementary fig 2

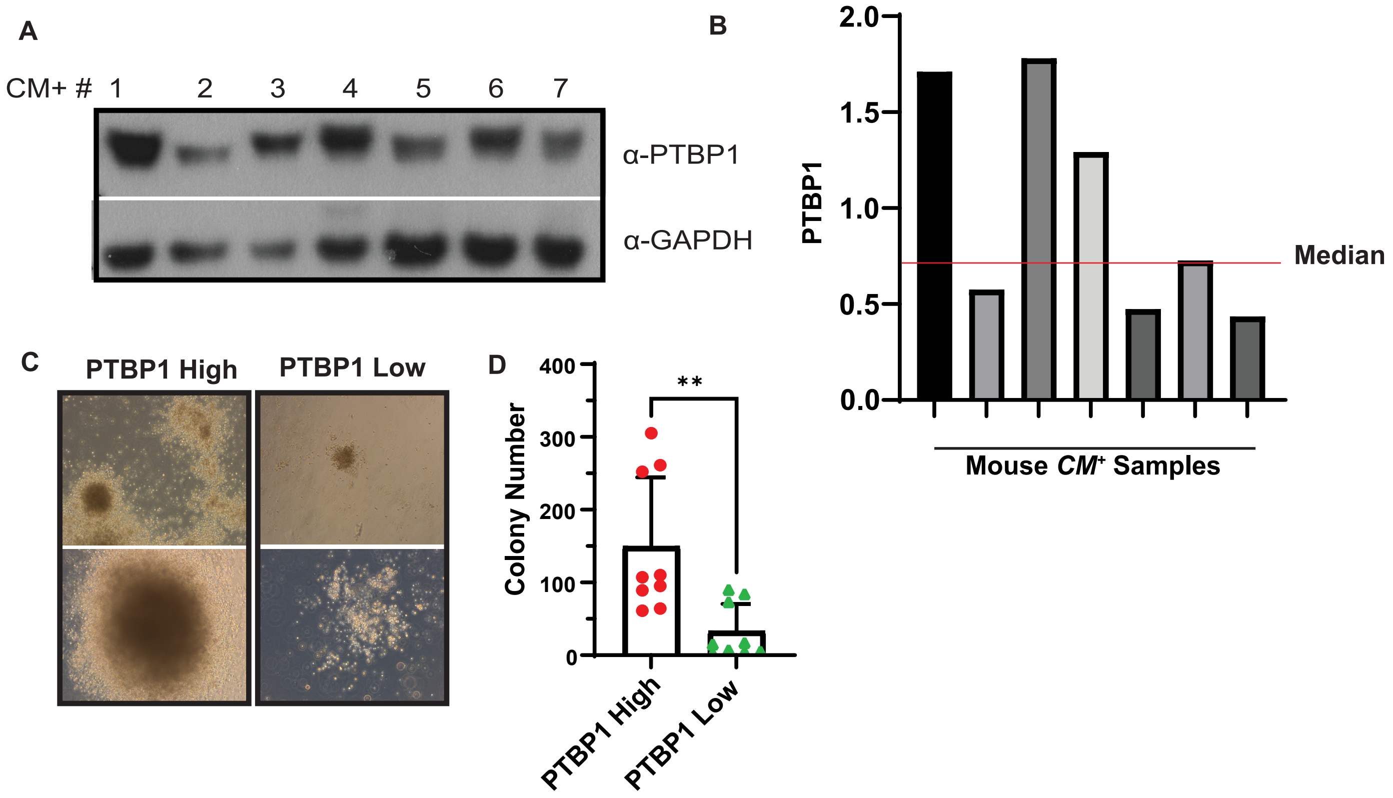

Supplementary fig 3

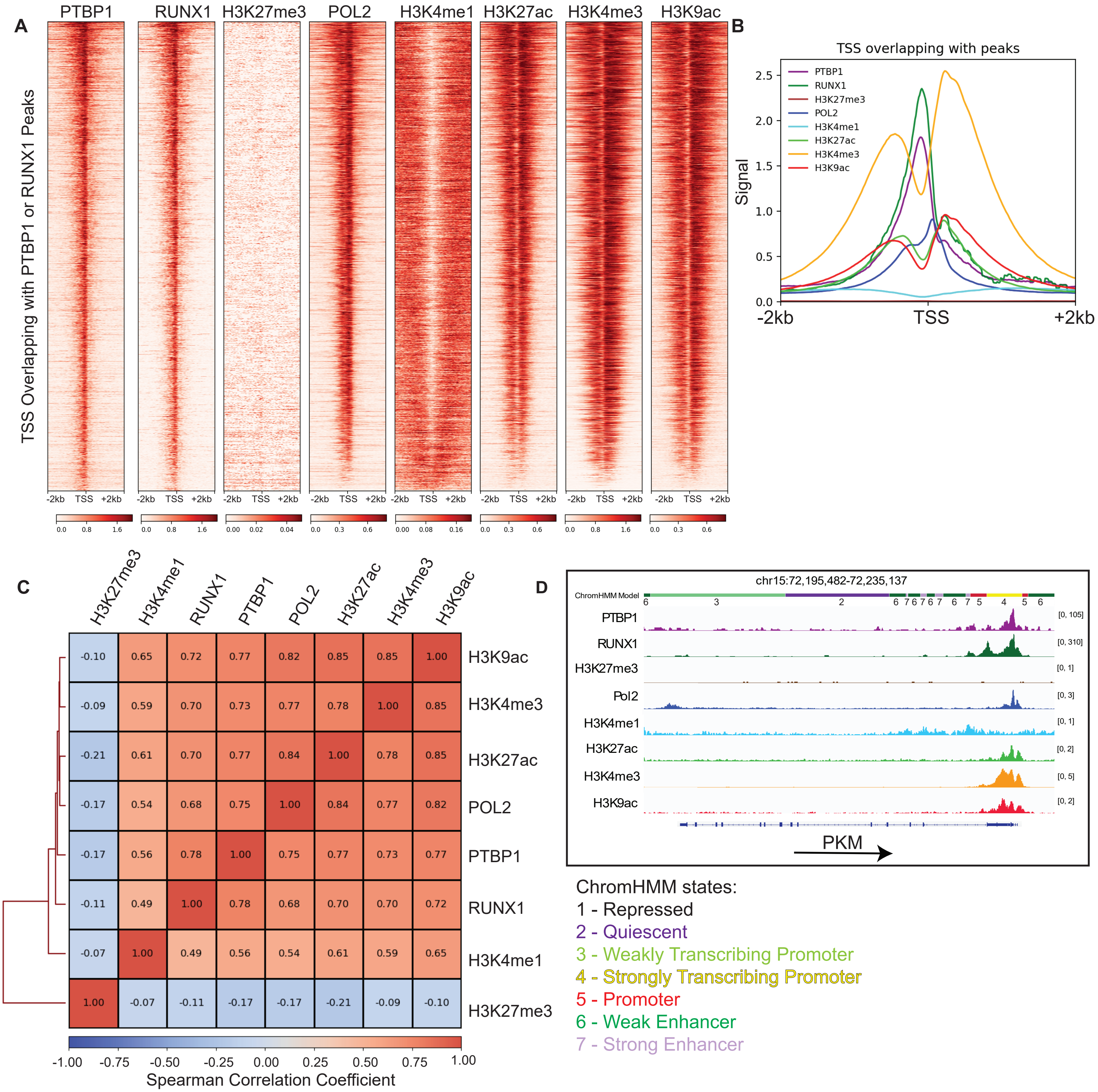

Supplementary fig 4

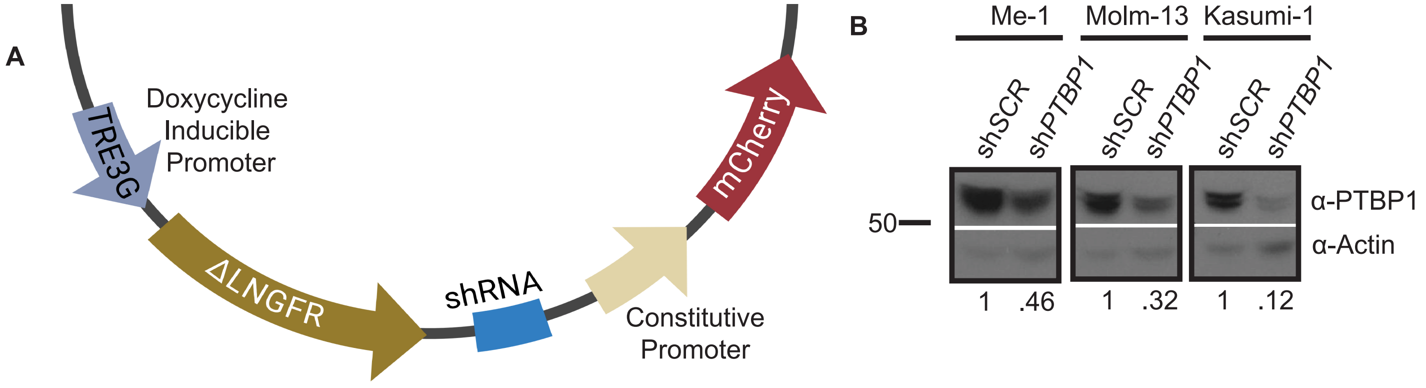

Supplementary fig 5

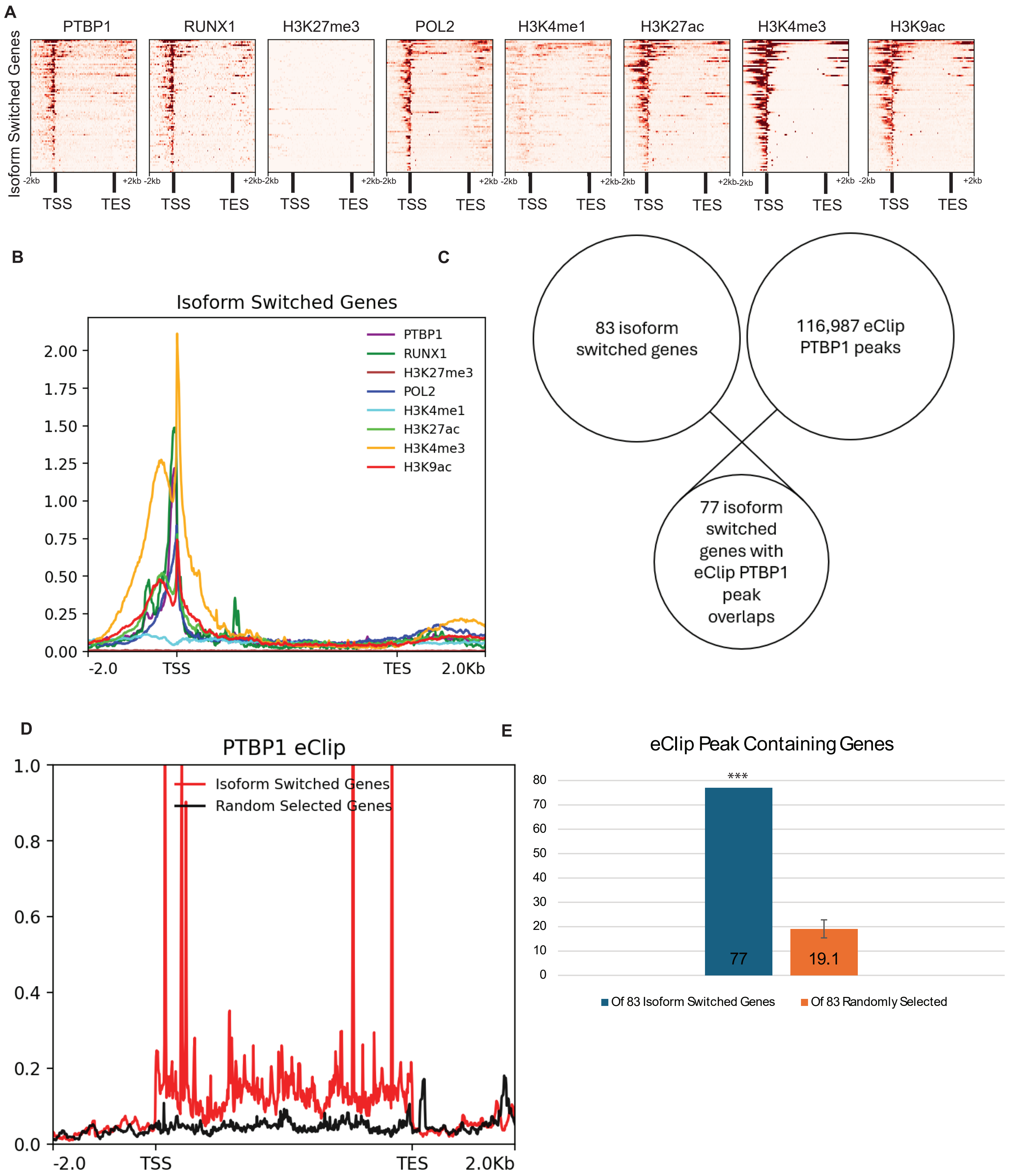

Supplementary fig 6

**A**

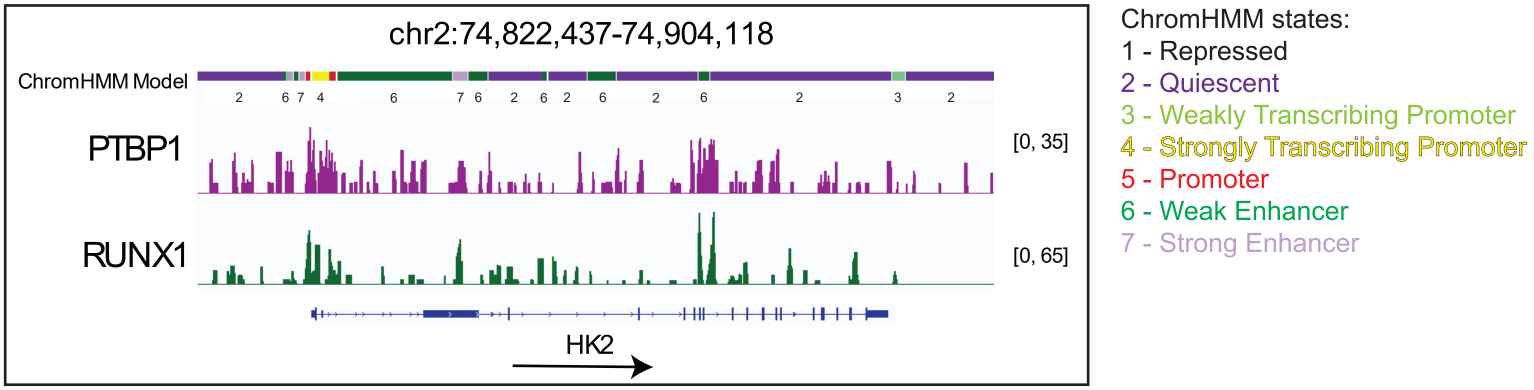


**B**

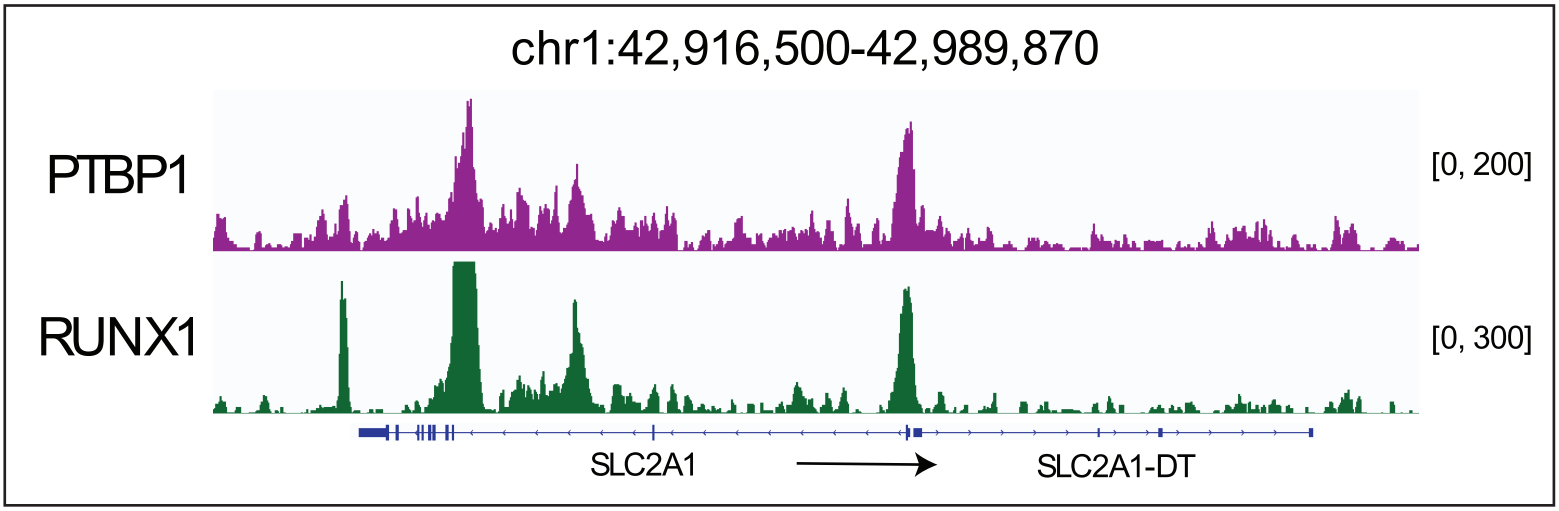
